# Cable-free brain imaging with miniature wireless microscopes

**DOI:** 10.1101/2022.06.20.496795

**Authors:** Yangzhen Wang, Zhongtian Ma, Wenzhao Li, Feng Su, Chong Wang, Wei Xiong, Changhui Li, Chen Zhang

## Abstract

The invention of the miniaturized microscope has enabled neuroscientists to investigate neural mechanisms in freely moving mice. A lot of efforts have been made to optimize performance of the miniaturized microscope. However, the tethered cables limit the ability of mini-microscope systems to record neural activity from multiple mice simultaneously. Here, we present a wireless mini-microscope (wScope) that enables both real-time remote control and data preview during animal behavior; this design also supports simultaneous recording from up to 8 mice. The wScope has a mass of 2.7 g and a maximum frame rate of 25 Hz at 750 μm by 450 μm field of view with 1.8 μm resolution. We validated the wScopes in video-recording of the cerebral blood flow (CBF) and the activity of neurons in the primary visual cortex (V1) of different mice. The wScope provides a powerful tool for brain imaging of free moving animals, including large primates, in their much larger spaces and more naturalistic environments.

## Introduction

Recording the activity of neurons in freely behaving rodents is a crucial way for neuroscientists to understand how neuronal networks process information in naturally and ethologically relevant behavior. Many electrophysiological techniques have been presented to record a large number of neurons, such as silicon probes and multi-tetrode arrays.(*1*) However, it is challenging to differentiate each neuron within the sampled population by the shapes of waveform. In 2011, the Schnitzer group developed the first wide-field miniaturized microscope for measuring neuronal activity; the device had a mass of 1.9 g, a lateral resolution of 2.5 µm, and a frame rate of 30 fps at 640 × 480 pixels(*1*). Later, the Silva group initiated the UCLA Miniscope project, featuring an open-source imaging platform with a mass of <3 g and a frame rate of 30 fps at 752 × 480 pixels.(*2*) With the help of the UCLA platform, researchers have invented a variety of additional miniature microscopes. For example, the Vaziri group used a microlens array and a constrained matrix factorization strategy to image a volume of 700 × 600 × 360 µm^3^ at a frame rate of 16 Hz and an axial resolution of 30 µm with a device weighing > 4 g.(*3*) Using a transparent polymer skull implant and a light-emitting diode (LED) array, the Kodandaramaiah group developed a whole-cortex miniscope with a field of view (FOV) of 8 × 10 mm, a mass of 3.8 g, and spatial resolution ranging from 39 to 56 µm.(*4*) These and other miniaturized microscopes have provided researchers with powerful tools to study the neural circuits of many sub regions of the brain, such as the cortex, hypothalamus, hippocampus, striatum and amygdala(Liberti et al., 2017; Kargl et al., 2020; Karigo et al., 2021; Kennedy et al., 2020; Kingsbury et al., 2019; Shin et al., 2020). However, the tethered cables used in these systems prevent multiple mice from freely moving together; thus, to study the complex behaviors of multiple mice in a single arena with more naturalistic conditions simultaneously, it would be ideal to develop a wireless microscope system that lacks these tethered cables, allowing total freedom of movement.

Among the many efforts made to accomplish this goal, wireless microscope systems have been reported to successfully record the activity of neurons in rodents(Supplementary table 1)(*11, 12*). The images from these microscopes are captured by a complementary metal-oxide-semiconductor (CMOS) imaging sensor and stored on a Micro SD card. While these systems enable mice to roam completely untethered during experiments, this comes at the expense of a lightweight design (3.8-4.5 g), the imaging resolution (⩽320 × 320 pixels), the frame rate (⩽20 fps) and the ability to preview data in real time. The weight of these microscopes is heavier than 10% of 8-week-old mice, and the behavior of animal could be influenced by microscope. In addition, the imaging parameters are also incomparable to the wired microscopes. In addition, the signals in SD microscopes are not transmitted in a truly wireless manner, and none of the microscope parameters can be adjusted during the experiments due to the lack of wireless communication between the user and the microscope. A wireless microscope with comparable parameters to traditional wired microscopes is needed to imaging the brain of multiple freely moving mice.

Here, based on the UCLA Miniscope platform, we developed a lightweight, wireless mini-microscope, named wScope, for brain imaging in freely behaving mice. Frequency Modulation (FM) technology is widely used in the fields of music broadcasting, TV signal transmission, satellite communications and cellular phone systems. We used FM technology to transmit high-quality image signals wirelessly at a performance level comparable to that of a wired microscope. Additionally, unlike previously reported wireless mini-microscopes, our system enables wireless control of many key parameters, including gain and LED power. Our system supports simultaneous brain imaging of up to 8 mice in the same arena. A lightweight, 100 mAh lithium battery provides sufficient power for over 15 minutes of imaging, and the camera can uniquely switch between standby mode and active mode through wireless control, which can maximize the efficiency of the camera and thus substantially extend the possible length of experiments by consuming energy to record videos only when needed.

To evaluate the image transmission quality of our wireless transmission system, we employed two parameters, namely, the peak signal-to-noise ratio (PSNR) and structural similarity (SSIM), to quantify the similarity between wireless and wired images(*13, 14*). We test the imaging and signal transmission performance of wScope, and recorded the cerebral blood flow (CBF) and neuronal activity of the primary visual cortex (V1) in different freely moving mice, both in an open space and in a tunnel, the latter of which is impossible with a tethered cable. The kinetics of vessel diameter change and neuronal calcium transients were recorded reliably by wScope, denoted a high performance of wScope for brain imaging. Our results demonstrate the power of our wireless microscope system as a tool to study CBF and neural circuits in multiple naturally behaving mice. The wScopes can help researchers study topics such as social behavior and cerebral disease in complex, custom-designed environments.

## Results

### Design of the wireless microscope system

The entire microscope has a volume of approximately 2.8 cm^3^ and a mass of 2.7 g, and it is powered by a lithium battery mounted on the back of the mouse. The frame rate of the wScope is 25 Hz, and the field of view is approximately 750 × 450 μm. The internal and circuit design of the microscope is shown in Fig. 1 a-b.

**Fig 1.**
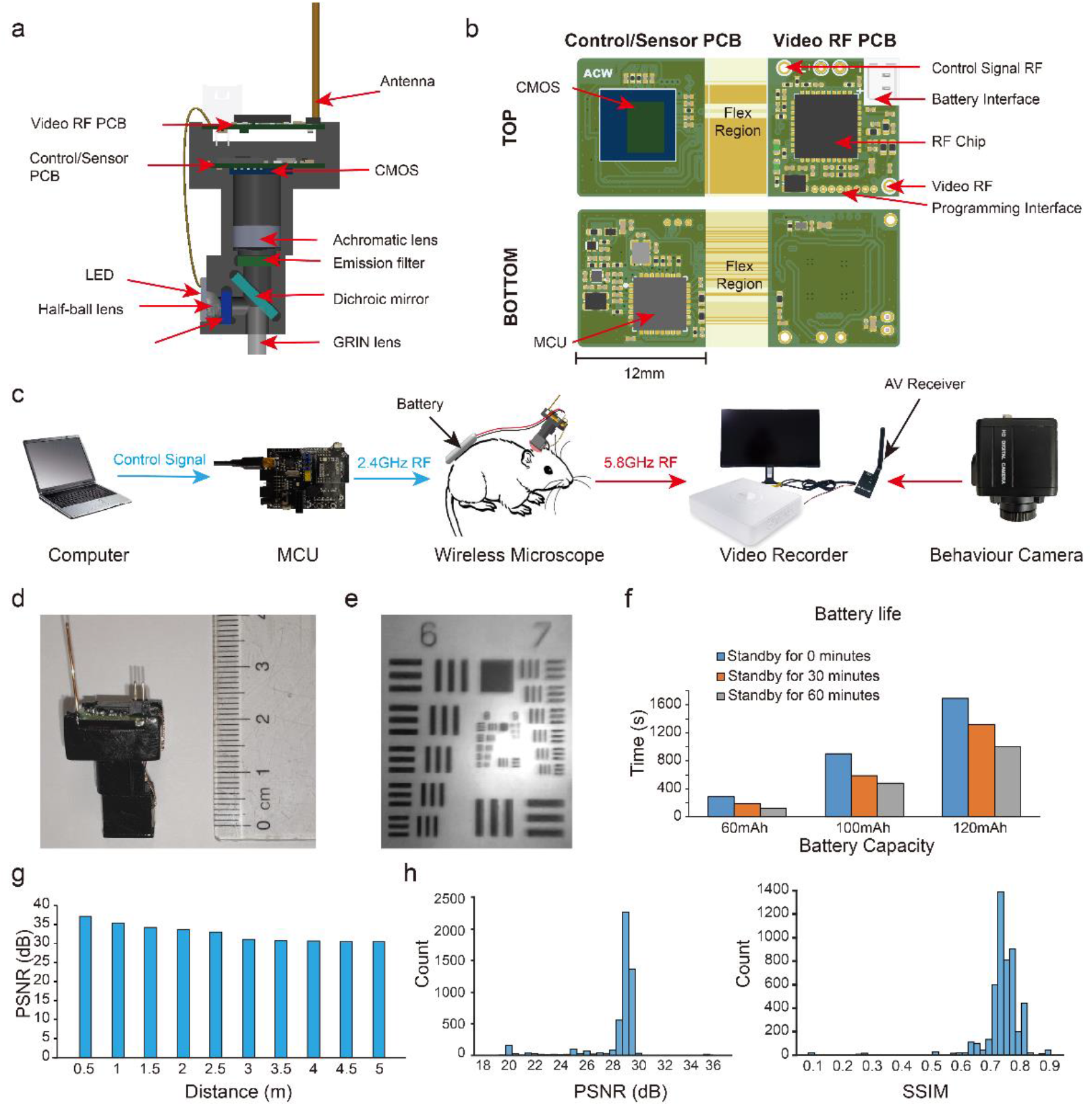
Wireless microscope imaging system. **a**, Design of the wireless microscope. **b**, Custom-built video RF and control/sensor PCBs, with views of the top and bottom components. **c**, Overview of the wireless microscopy system. **d**, Image of an assembled wireless microscope. **e**, Image taken by the wireless microscope of a USAF1951 Resolution Test Target. **f**, Battery usage time and usage time after standby. **g**, PSNR at different distances. **h**, Histogram of per frame PSNR and SSIM of the wireless conditions, as compared to the wired condition (at 2 meters).

The printed circuit board (PCB), shown in Fig. 1 b, contains two parts connected by a flexible connector. Once folded and placed, the circuit board consists of upper and lower layers. The upper circuit board includes a radio frequency (RF) chip; an antenna contacts for receiving control signals and transmitting video signals; and interfaces for power, LED. The main components of the lower circuit board are a CMOS image sensor and a 2.4 GHz RF transceiver with an embedded microcontroller unit (MCU). This folded design not only reduces the amount of space needed but also facilitates heat dissipation to prevent overheating. With these improvements, no significant behavioral difference was observed in mice when they wore the wScope or not.

Frames from the image sensor are sent through a 5.8 GHz radio frequency signal to an audio/video (AV) receiver and then viewed and saved in real time using a video recorder. We can control various elements for multiple microscopes separately, including the microscope CMOS gain, the brightness of the excitation LED and the microscope standby function.

### Test of the imaging and signal transmission performance of the wireless system

We measured the continuous working time of batteries of different sizes and capacities (Supplementary table 2). Additionally, to verify the effectiveness of the standby function in conserving battery life, we also measured the working time after half an hour and one hour of standby (Figure 1 f). As expected, larger capacity batteries demonstrated increasing battery life, which decreased for longer standby durations, we chose a lithium battery with a capacity of 100 mAh, a mass of 1.7 g, and a size of 4.0 mm × 10 mm × 25 mm for the in vivo experiment.

We also tested the image quality as a function of the source-receiver distance, as shown in Fig. 1 g. Taking the image at 0 distance as the reference image, we took images at 0.5 m intervals from 0.0 to 5.0 m, and the calculated PSNR values were all above 30. Thus, we considered our equipment to have almost constant transmission quality within a distance of 5 m. The PSNR value of the image received after wireless transmission was approximately 28∼30 dB, and the SSIM value was approximately 0.7∼0.8 (Figure 1 h), which demonstrates that our wireless transmission can guarantee reliable image quality(*13–15*).

### Imaging of CBF in active mice

To evaluate the performance of the new system for in vivo brain imaging, we built an arena and placed a camera above it to monitor the behavior of the animals. Two sides of the arena were made of transparent acrylic plates, allowing the animals to see visual stimuli presented on screens outside the transparent walls (Fig. 2 a). The microscope and battery, connected by thin electrical wires, were mounted on the skull and back of the mouse, respectively. Before the experiments, the mice underwent adaptation training with a dummy microscope mounted on their heads; after training, the mice showed robust locomotor behavior (Supplementary figure 1). The cerebral microcirculation of V1 in freely behaving mice was labeled with fluorescent dye, and the hemodynamic responses to visual stimuli were analyzed to evaluate the CBF (Fig. 2 b). Concurrently with brain imaging, the behavior of the mice were recorded, and a visual stimulus was presented when the mice turned toward the screen. After the presentation of stimuli, the vessels in V1 quickly began to dilate (0.45 s ± 0.11) and reached their peak dilation amplitude (19.46% ± 4.21) in 1.66 s ± 0.10, followed by contraction in 5.69 s ± 0.36 to reach their peak contraction amplitude (7.70% ± 1.57), after which they gradually returned to baseline (Fig. 2 c-e). The vascular kinetics were correlated with visual stimulation, and the amplitude of the vessel dilations was significantly higher than that of the contractions. These results demonstrate that our wireless microscope system can reliably record the changes in cerebral vessels in freely moving mice, enabling brain hemodynamic imaging studies for a variety of topics, such as cerebrovascular disorders and neurovascular kinetics.

**Fig 2.**
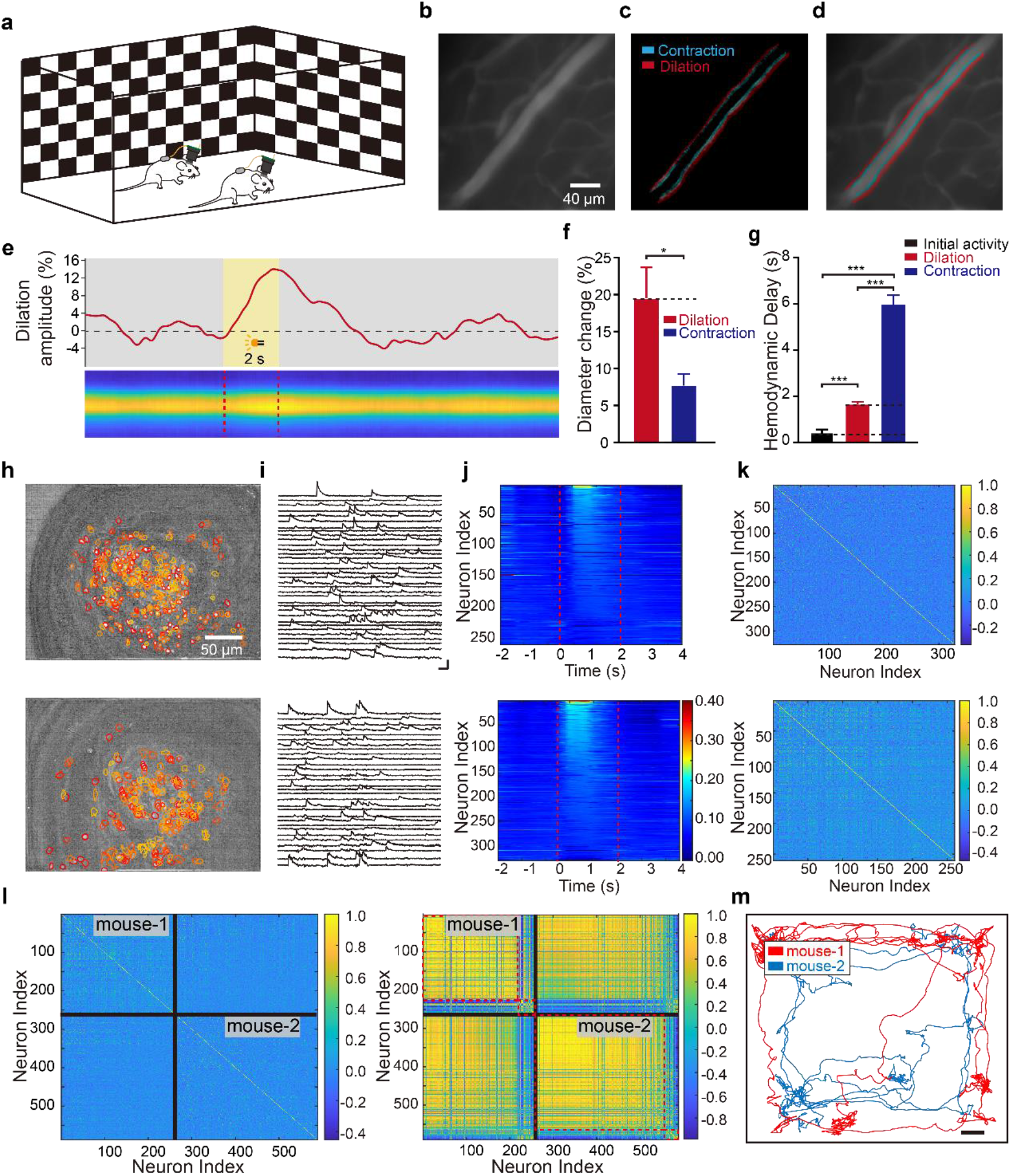
In vivo brain imaging in freely moving mice. **a**, Illustration of mice fitted with the wireless microscope system in an open arena. **b** Example image of brain vessel imaging using the wireless microscope system. **c**, Changes in the diameter of the vessel during visual stimulation. **d**, Merged image showing the vessel and the diameter change. **e**, Example trace (c1) and line scan heatmap (c2) of vessel diameter change during stimulus presentation. **f**, The absolute diameter changes for vessel dilation and contraction during visual stimulation. **g**, The delays of the initial phase, the peak of the dilation phase and the peak of the contraction phase of vessels with respect to the onset of visual stimulation. **h**, Signal-to-noise ratio maps and the location of each neuron in the V1 of the two mice. **i**, Representative transient calcium traces from the V1 neurons of two active mice in the dark phase. **j**, Heatmap of neuronal calcium transients during visual stimulation. The red dashed lines indicate the stimulus phase. **k**, Matrix of the Pearson correlation coefficients between the calcium transients of pairs of neurons during the dark phase. **l**, Matrix of the Pearson correlation coefficients between the calcium transients of different neurons from the two mice during the dark phase (left) and the stimulus phase (right). The black line separates the neurons of two mice from each other; the red dashed lines group highly correlated neural populations. **m**, Locomotion trajectories of two mice fitted with the wireless microscope system within the arena (Scale bar: 5 cm).

### Recording the neuronal activation of the primary visual cortex in two mice simultaneously

We next tested the performance of the system in simultaneously recording neuronal activity from two mice. Four weeks after the injection of rAAV2/9-hsyn-GCamp6m in V1, the mice were trained using dummy microscopes for one week. During the experiments, the two mice were allowed to freely explore and engage in social interaction in the arena. The arena and visual stimuli were the same as those in Fig. 2 a. The population neuronal activity in V1 was analyzed using extended constrained nonnegative matrix factorization (CNMF-E) and ImageCN(*16–18*); 260 spatial components were detected in one of the mice and 320 in the other (Fig. 2 f). The temporal components of neuronal activities were also extracted and calcium signals were distinguishable from the background noise (Fig. 2 g). The neural population in the V1 areas of both mice demonstrated strong activity beginning ∼0.2 s after the onset of stimulus presentation (Fig. 2 h). To evaluate the performance of the wireless system in capturing the two recordings independently and simultaneously and confirm the absence of interference between them, we used the absolute value of the Pearson correlation coefficient, a statistic for measuring the linear relationship between each pair of neurons, to quantify the inter- and intragroup correlations of the neuron populations. In the dark phase (no light in the arena) of the experiments, no visual stimulus was presented, and both mice showed weak correlations between the activities of different pairs of neurons within the population (Fig. 2 i). Moreover, the population correlation between the two mice was low during the dark phase, the signal received by two mice were independent, indicating that no interference was detected during simultaneous recording with two microscopes (Fig. 2 j, left). However, during visual stimulus presentation, the neuron populations tended to fire simultaneously. Thus, the correlation of activity between pairs of neurons in both mice was higher in this phase than during the dark phase, when these populations were quiescent (Fig. 2 j, right). The behavior of the animals was recorded with a camera; the corresponding movement trajectories of the two mice are shown in Fig. 2k. Overall, we recorded the calcium transients of V1 neurons in two mice, and no signal interference was observed between different channels of microscopes, indicate that our system provides a powerful tool for recording the activity of neural populations in multiple freely moving mice with excellent spatial and temporal resolution.

## Discussion

In this paper, we developed a wireless miniaturized microscope system for simultaneous brain imaging in multiple mice. Using FM technology, wireless signal transmission and system control were achieved.

wScope has similar imaging performance to existing wired microscopes, while its wireless hardware allows the experimenter to study the behavior of multiple freely moving mice at once; it can even be used while the animals are in enclosed space (Supplementary figure 2). Our system has a smaller mass (2.7 g), higher resolution (640 × 480 pixels) and a faster frame rate (25 Hz) than other recently reported wireless microscope systems. Moreover, we added the ability to remotely control imaging parameters and switch the device between standby mode and recording mode to improve the imaging performance of our system and extend its battery life. As shown by our data, this wireless system achieved excellent signal transmission performance, and chronologically ordered images of CBF and neuronal activity in V1 were recorded with high fidelity and resolution. Using wScopes, we successfully imaged 4 mice simultaneously in the same arena (Supplementary figure 3-4). We further compared these data with that from previously published literature on cerebral hemodynamic changes in response to stimuli; our data were comparable to those in the literature, demonstrating the reliability of our system(*19–21*).

In the current version of the microscope, wireless transmission was realized by the use of analog signal frequency modulation. This scheme has poor resistance to interference and is not very sensitive to weak signal changes and thus needs to be tested in a relatively clean electromagnetic environment. Furthermore, digital transmission schemes that are currently in common use, such as Wi-Fi and Bluetooth, generally cannot provide sufficient transmission rates (for example, if a frame rate of >20 Hz is needed, then the transmission speed needs to be 70 Mbps or more) or keep power consumption sufficiently low. Therefore, in future research, we will consider combining real-time data compression techniques with new digital transmission schemes to design a new generation of miniature wireless microscopes. With the advancement of semiconductor technology and chip design, digital wireless transmission solutions with ultralow power consumption are possible.

The emergence of new materials will promote the development of new battery technology, which will extend the life of the system. The advancement of wireless charging technology will further reduce the weight that the mice must carry, even making it possible to eliminate battery-life limitations completely and monitor neuronal activity in the long term, e.g., days or months. These capabilities will create exciting new experimental opportunities.

Miniaturized microscopes play an important role in neuroscience, helping researchers study neural circuits in freely behaving animals. Our wireless microscope system could extend the range of applications for brain imaging in freely moving rodents, including some animal experiments are conducted in closed tubes or boxes, which would not permit the use of wired microscopes. Furthermore, the wScope can be used in free-moving larger animals, such as large primates, that cannot use the tethered cable. Even more, with a larger brain size and body weight, multiple wScopes can be mounted on one head that allows the study of cross-brain neuron connections in free-moving large primates.

## Methods

### Wireless system design

The goal of our study was to observe the brain nerve activity of multiple freely moving mice at the same time, and to observe the image in real time. Our proposed system is based on the open-source, mini-microscope system developed at UCLA; specifically, we began with the physical shell of the miniscope and introduced our own modifications.

Each microscope has a blue LED with a spectral peak of ∼475 nm (LXML-PB01-0040), which is supplied with energy by an LED interface on the circuit board. The excitation light generated by the LED passes through a half-ball lens for collimation and is then passed through an excitation filter (Edmund Optics, ET470/40x, 3.5 × 4 × 1 mm^3^) and reflected using a dichroic mirror (Edmund Optics, T495lpxr, 4 × 6 × 1 mm^3^). The emitted light leaves the objective GRIN lens in a substantially parallel direction, passes through a dichroic mirror and then passes through an emission filter (Edmund Optics, ET525/50m, 4 × 4 × 1 mm^3^). Finally, an achromatic lens (Edmund Optics, 5 mm diameter, 15 mm FL) is used to focus the light onto a CMOS imaging sensor (OV7960).

Our system uses nRF24LE1 chips, a 2.4 GHz RF transceiver with embedded microcontroller, to realize wireless control. The nRF24LE1 is mounted onto the microscope circuit board and is used to wirelessly receive instructions to control the microscope parameters. Each microscope is assigned a unique number so that its parameters can be controlled separately during the experiment without affecting other microscopes. Another nRF24LE1 is installed on a development board and connected to the computer through a serial port. The parameters that need to be controlled are selected on the computer (Supplementary figure 5) and are then sent to the development board through the serial port. The development board sends the instructions as a 2.4 GHz radio-frequency signal.

The OV7960 image sensor generates streaming pixel data at a constant frame rate with analog output. The images are wirelessly transmitted using an RTC6705, a 5.8 GHz-band FM transmitter.

To make efficient use of power, the power consumption of the RTC6705 is set to +2 dBm. The RTC6705 has 24 fixed channels in the 5.8 GHz band. A fixed channel can be assigned to each microscope through the embedded program of the nRF24LE1 so that multiple microscopes can be used at the same time without interfering with each other.

### System performance testing

In order to test the quality of the wireless transmission, we use the wScope to image the fluorescence micrometer and sent and analyzed video signals via both wired and wireless transmission to evaluate the quality of the images obtained in real time. For this purpose, we connected the CMOS image signal directly to the video recorder through a video buffer module (AD8079) and compared it with the wirelessly transmitted image signal. We chose a transmission distance of 2 m, which is sufficient for most of our experimental needs.

The PSNR and SSIM were chosen to quantify the quality of the transmitted images and the similarity between the two images, respectively. To calculate the PSNR, we must first calculate the mean squared error (MSE). For two *m* × *n* images x(the wired image) and y(the wireless image), the MSE is defined as:

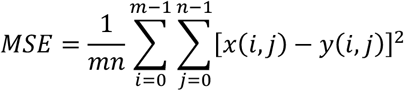

The PSNR is calculated from the MSE with the following formula:

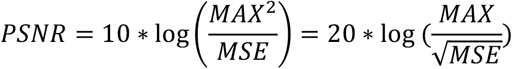

MAX is the maximum value of the color scale that is applied to each point in the image. If each sampling point is represented by 8 bits, then MAX is 255. The smaller the MSE is, the larger the PSNR; thus, the larger the PSNR is, the better the image quality.

Continue the previous two images x and y, the SSIM between them can be calculated as follows:

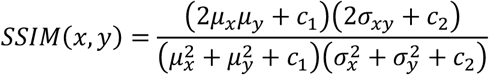

*µ*_*x*_ is the average of image x, and *µ*_*y*_ is the average of image y. 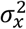 is the variance of 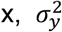 is the variance of y, and 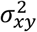 is the covariance of x and y. *c*_1_ = (*k*_1_ *L*)^2^, *c*_2_ = (*k*_2_*L*)^2^, *k*_1_ = 0.01, and *k*_2_ = 0.03. The larger the SSIM value, the more similar the two images are; when the two images are exactly the same, the SSIM is equal to 1.

To test the wireless transmission performance of our system, we measured the received image signals at different distances. Taking the image with a transmission distance of 0 as a reference, the PSNR value was calculated between the image signal received at each distance and the reference image signal.

### Animal surgery

All animal experiments were conducted at Capital Medical University Laboratory Animal Center, an Association for Assessment and Accreditation of Laboratory Animal Care (AAALAC)-approved animal facility. Wild-type C57BL/6 mice (male, aged 8–12 weeks) were purchased from Vital River Laboratories (Beijing, China) and maintained on a 12/12-hour reversed dark-light cycle. All experiments were performed during the light cycle. Before surgery, the mice were anesthetized with tribromoethanol (240 mg/kg, Sigma), and a craniotomy was performed over the primary visual cortex of the right hemisphere (coordinates with respect to bregma: anteroposterior, -2.8 mm; mediolateral, 2.5 mm; dorsoventral, 0.2 mm). After the craniotomy, a 1.8 mm diameter GRIN lens (Edmund Optics) was placed on the surface of the cortex without damaging the tissue. The gap between the GRIN lens and the skull was covered with Kwik-Sil (World Precision Instruments) to protect the cortex. A metal baseplate was mounted on the skull using cyanoacrylate adhesive and dental acrylic. After surgery, the animals were allowed to recover for 1 week, during which they were intraperitoneally injected with ceftriaxone sodium (200 mg/kg) and dexamethasone (5 mg/kg) every day to prevent inflammation and edema. One week after the surgery, the mice mounted with a 3D printed dummy microscope on their heads, for robusting the locomotor behavior.(*22*)

### Cerebral vessel imaging in awake mice

Before imaging, the animals were anesthetized with isoflurane. Fluorescein isothiocyanate-dextran (2% w/v, MW 70,000, 50 mL, Sigma) was administered into the femoral vein to label the blood plasma. A microscope was mounted on the mouse’s head, and the focal plane was adjusted manually to show a clear image of the vessels. One piece of self-gripping fastener material was taped to the battery, and the complementary piece was sutured to the back of the mouse. Subsequently, the microscope and battery were connected with a wire. The gain and LED power were adjusted to the proper values (vessel is clearly visible, and no over exposure), and the device was powered down immediately to conserve battery life. All these adjustments were made by remote control. Still anesthetized, the animal was placed in the arena, a 50 × 50 cm open field made by polymethyl methacrylate; 10 minutes after waking, imaging was started. The behavior of the mouse was observed by the experimenters, and stimuli were presented for 2 s when the animal’s head was turned toward one of the screens placed behind two transparent walls of the arena.

### Virus infusion for calcium imaging

The location of V1 was determined using stereotactic coordinates. After craniotomy, the dura was removed around the injection area. A virus (AAV2/9.syn.GCaMP6m.WPRE.SV40, diluted to a titer of 4 × 10^12^ vg/mL) was drawn into a prepared glass pipette (tip size: 0.3-0.5 mm) through negative pressure. To create a seal against the surface of the brain, the glass pipette was slowly lowered again by approximately 450 μm to press on the pia without breaking through it. Viral infusion was started after 2 min and proceeded at 60 nL/min for 10 min(*23*). The craniotomy was covered with a small piece of glass coverslip after infusion. The animals recovered for 7 days after the craniotomy and were intraperitoneally administered ceftriaxone sodium (200 mg/kg) every day to prevent inflammation.

### Data processing and analysis

For CBF analysis, the changes in the diameter of cerebral blood vessels were analyzed. All vessels with clear boundaries in the recorded video were selected by the experimenter. For each vessel, a line was manually drawn across the vessel, perpendicular to its axis. The one-dimensional array of gray values of each pixel along the line was used to calculate the diameter of the vessel. The maximum value in the array was calculated. On the left and right sides of the index corresponding to this maximum value, the five lowest gray values were found, and their average values were calculated as left and right references. One-half the mean of the maximum value and the left/right reference value was calculated as the left/right threshold value. For the left and right sides, the linear index of the maximum element that was less than the left/right threshold value was calculated, and the distance between the left and right sides was denoted as the diameter of the vessel. Since the images recorded by the microscope were arranged in chronological order, the activity of the vessel could be illustrated as a trace of its diameter from image to image.

For neuronal activity analysis, CNMF-E was used to detect the spatiotemporal components of activity in neuronal populations. To quantify the correlation between each pair of neurons, the Pearson correlation coefficient (R) was calculated as follows:

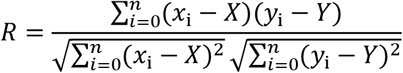

where *x*_i_ and *y*_i_ are the normalized values of the calcium transient traces, X and Y are the mean values of the traces, and n is the trace length.

## Supporting information

multi-channel recording demo video

Video of a mouse moved in the tunnel

## Acknowledgments

This work was supported by grants from the National Basic Research Program of China [2017YFA0105201]; the National Science Foundation of China [81925011]; Key-Area Research and Development Program of Guangdong Province [2019B030335001]; The Youth Beijing Scholars Program (015), Support Project of High-level Teachers in Beijing Municipal Universities [CIT&TCD20190334]; and Beijing NSF Program and Scientific Research Key Program of Beijing Municipal Commission of Education [KZ201910025025]; The National Key Research and Development Program from the Ministry of Science and Technology of the People’s Republic of China [No. 2017YFE0104200]; and the National Natural Science Foundation of China [No. 81421004].

## Author contributions

C.Z., and C.L. conceived and planned the experiments. C.L, C.Z., W.L., Z.M. and Y.W. designed and built the system. Y.W. and Z.M. carried out the experiments. Y.W., Z.M., F.S. and C.W. processed the experimental data, performed the analysis. All authors were involved in discussing the results, writing the manuscript and had approval of the final versions.

