## Supplementary material for "Cable-free brain imaging with miniature wireless microscopes": Supplymentary information

**Supplementary table 1 Comparison of microscope parameters with SD card storage solution**

| | wScope | Schnitzer group's microscope | SD card solution 1 | SD card solution 2 |
|---|---|---|---|---|
| **Weight** | 2.7g | 1.9 g | 3.9g | 4.5g |
| **Resolution** | 640*480 | 640*480 | 200*200 | 320*320 |
| **Frame Rate** | 25 Hz | 36 Hz | 10 Hz | 20 Hz |
| **Battery Life (100 mAh)** | 16min | \ | 16min | 30min |
| **Field of View** | 700*450μm | 600*800μm | 500*500μm | 700*450 |

**Supplementary table 2. Battery Information**

| Size | Capacity | Weight |
|---|---|---|
| 4.0mm*10mm*15mm | 60mAh | 1.32g |
| 4.0mm*10mm*25mm | 100mAh | 1.7g |
| 6.0mm*12mm*25mm | 130mAh | 3.3g |

**Supplementary figure 1. Animal recovery training using dummy microscope after surgery**

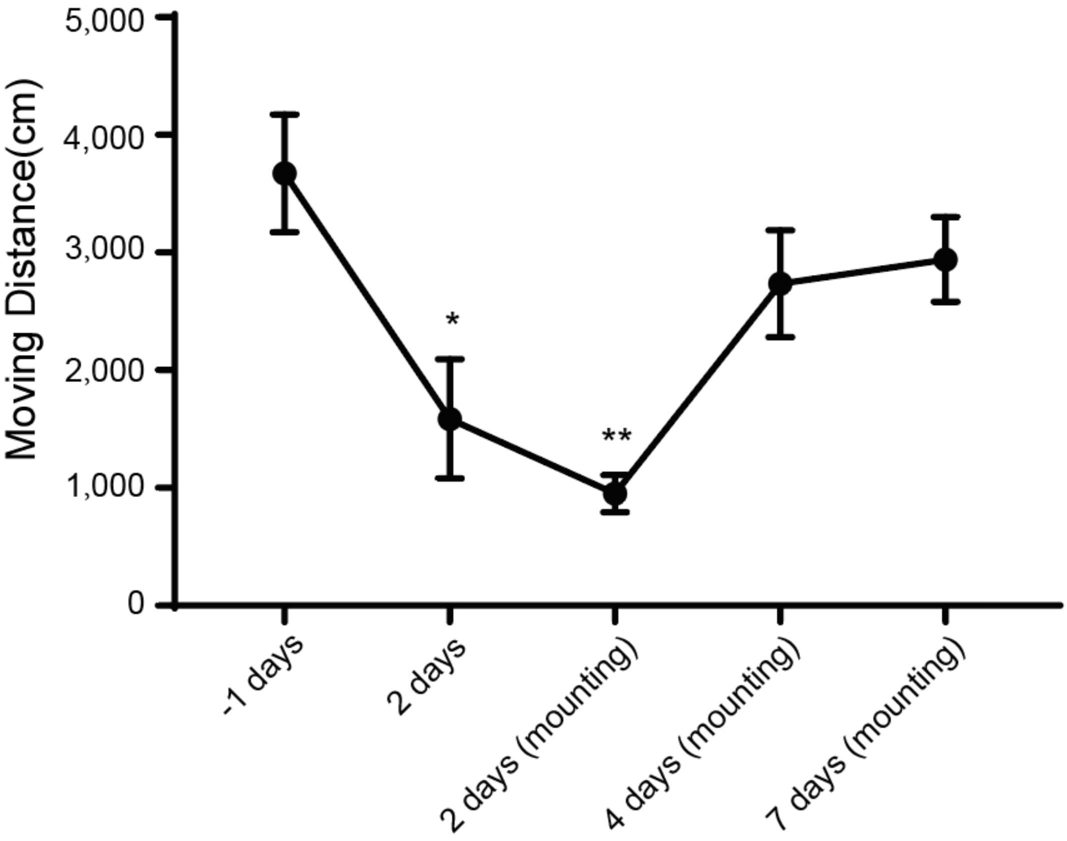


Animal moving distance in arena for 10 minutes was calculated to evaluate the effects of the surgery and the mounting of microscope to mice. 1 day before surgery, the moving distances were calculated as control. 2 days after surgery, the moving distances were decreased significantly, and a further decreased was observed after the mounting of dummy microscope. After 2 days training, the behavior of mice was recovered.

**Supplementary figure 2. Evaluation imaging PSNR while the animal was in enclosed space**

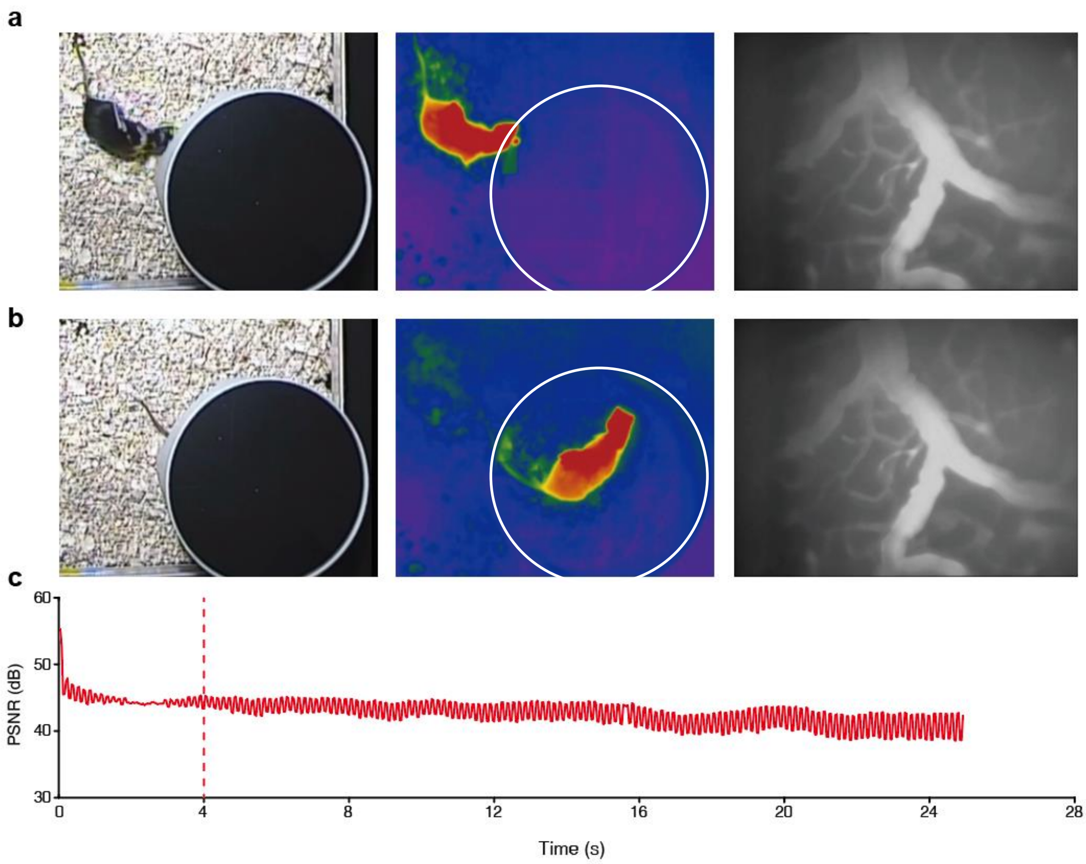


**a**. Mouse mounted with wScope was exploring the door of the enclosed space. Left: visible light behavior camera imaging, middle: thermal infrared behavior camera imaging, right: vessel imaging using wScope; **b**. Same depiction as a, except the body of mouse was in inside the enclosed space; **c**. PSNR of vessel images before and after the mouse entered the enclosed space, as compared to the first frame, red dash line denotes the moment of mouse's head is entering the space.

**Supplementary figure 3. Simultaneously 4 mice locomotion recording in the same arena**

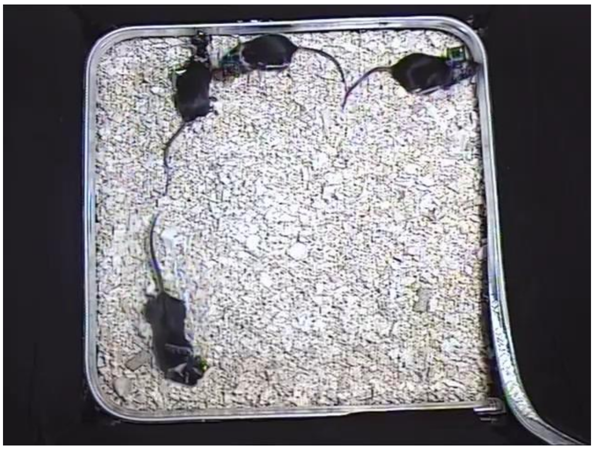

4 mice mounted with wScopes were freely locomoting in the same arena simultaneously.

**Supplementary figure 4. Cerebral vessel imaging in 4 mice simultaneously**

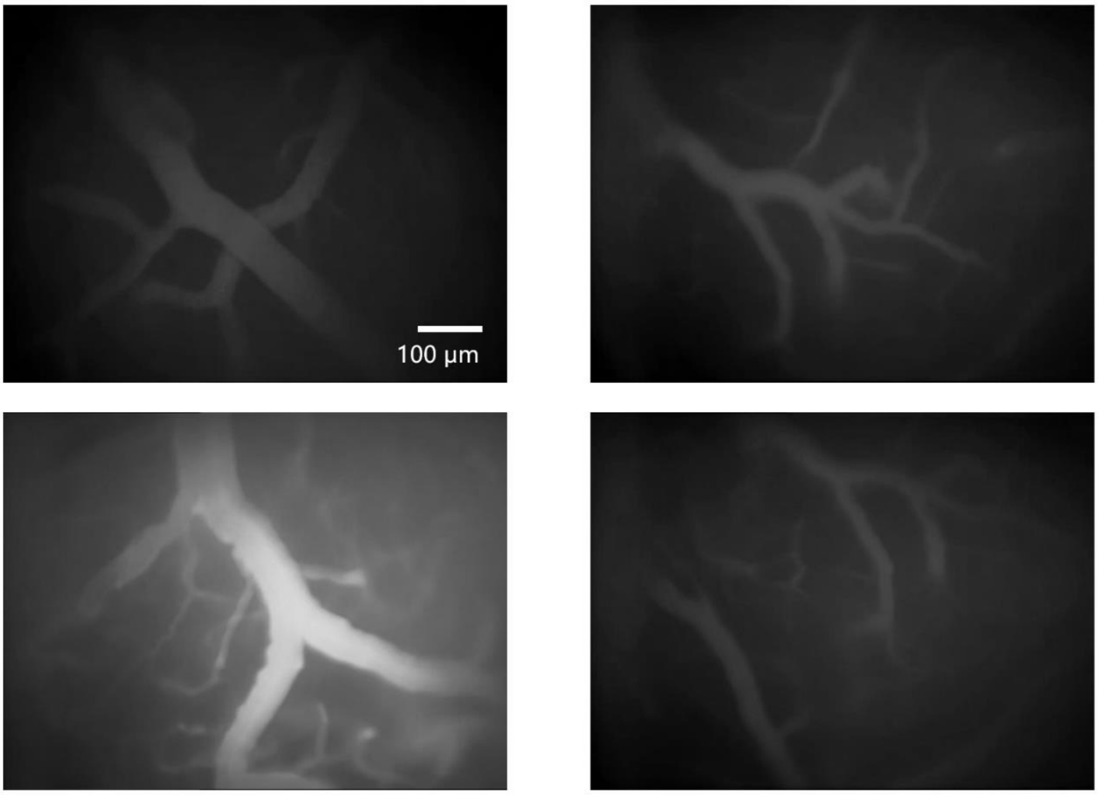


4 mice brain vessels imaging simultaneously using wScope.

**Supplementary figure 5. Wireless microscope control program user interface**

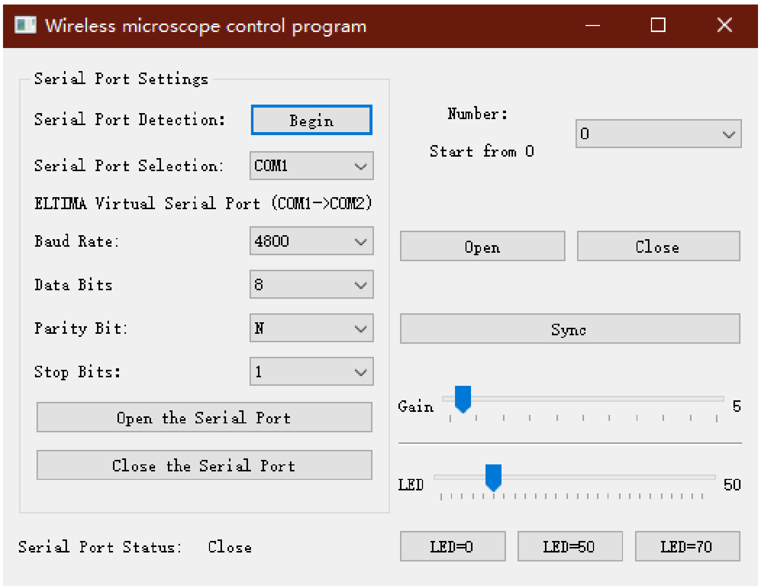
